# Self/non-self discrimination in tendrils of the vine Cayratia japonica with electrical signals

**DOI:** 10.1101/2025.07.31.667820

**Authors:** Takahiro Homma, Yoshio Kami

**Affiliations:** Center for Industrial and Governmental Relations, University of Electro-Communications, 1-5-1 Chofugaoka, Chofu, Tokyo 182-8585 Japan; Professor Emeritus, University of Electro-Communications

**Keywords:** Cayratia japonica, electrical signal, tendril coiling, self/non-self discrimination

## Abstract

Recent studies have shown that the vine Cayratia japonica (C. japonica) has a self/non-self discrimination ability via its tendrils. However, the mechanism of self/non-self discrimination in tendrils of C. japonica remains unclear. Here, we show that tendrils of C. japonica discriminated between self and non-self plants with electrical signals generated by the contact of tendrils. We conducted touch experiments with C. japonica and detected electrical signals generated by the contact of tendrils. We found that when the tendril contacted the stem and did not coil around it, the electrical signal was transmitted from the stem to the tendril. On the other hand, when the tendril coiled around the stem, no electrical signal was transmitted. These results suggest that the electrical signals inhibited the coiling of tendrils, and the following experiment was conducted to verify this hypothesis: (1) when the tendrils did not coil around the stems after contacting the stems, the tendrils were removed from the stems and contacted a stick to coil around it. (2) Contact stimuli were applied to the part of the stems where the tendrils did not coil around, and the tendril uncoiled. We found that self/non-self discrimination was realised by electrical signals that inhibited coiling generated by contact stimuli.

## 1. Introduction

Tendril movements have long fascinated biologists, dating back to Charles Darwin’s studies on climbing plants [1]. A tendril circumnutates and searches for a potential support. When the tendril touches the support, it stops circumnutating and begins coiling [2]. Previous studies have shown that all higher plants may use electrical signals to regulate various physiological functions [3]. Tendrils stop their autonomous circumnutations immediately due to the loss of action potentials, a type of electrical signal, under diethyl ether anaesthesia [4]. Additionally, non-damaging stimuli (e.g., cold, contact stimuli) evoke the propagation of action potentials throughout the plant body [3, 5, 6]. The contact stimuli generate action potentials in tendrils because the coiling of the tendril is a response to contact stimuli [7, 8]. The contact stimuli further trigger a complex chain of events, including fast ionic processes as well as chemical signalling to coordinate the reactions of the whole organ [9]. Several models for such chains were proposed as follows:

1. Contact stimulus -> action potential -> increase in free auxin -> asymmetrical response on two sides of the tendril [10],

2. Contact stimulus -> (action potential) -> ethylene production -> coiling [2].

Other previous studies have also shown that an increase of transmembrane proton and calcium fluxes, in addition to the involvement of ATP and light, is related to the beginning of the coiling movement in tendrils [11]. However, the full extent of the coiling mechanism remains unknown.

Electrical signals include variation potentials (VPs) or slow wave potentials as well as action potentials (APs). While APs are generally evoked by non-invasive stimuli, VPs are triggered mainly by wounding. APs and VPs are transmitted via the phloem over long distances and via plasmodesmata over short distances from cell to cell. APs propagate with velocities of 0.5–20 cm/sec, whereas the velocity of VPs is in the range of 0.1–1.0 cm/s [12]. VPs vary with the intensity of the stimulus [13].

Animals’ ability to distinguish between self and non-self is crucial because, without it, they may lose body parts and, in the worst case, die. In humans, after experiencing hand regard, which is the eye-hand coordination of an infant observed during the early months of their development, an infant may recognise their own hands [14, 15]. There has been some evidence that plants also have distinctions between self and non-self. Darwin reported that two tendrils, each growing from a nearby shoot of one Echinocystis, are repeatedly drawn across one another, but they do not once catch each other [1]. Tendrils of the perennial vine Cayratia japonica (C. japonica) are more likely to coil around stems of physiologically severed self plants than those of physiologically connected self plants; that is, self/non-self discrimination in tendrils is mediated by physiological connection [16]. In addition to the tendrils of the C. japonica, tendrils of Momordica charantia var. pavel (Cucurbitaceae) and Passiflora caerulea (Passifloraceae) can discriminate between self and non-self plants [17]. Plants can also discriminate between self and non-self plants via roots [18-21]. The tendrils of C. japonica avoid coiling around a conspecific leaf based on contact chemoreception for oxalate compounds [22]. However, the mechanism of self/non-self discrimination in tendrils of C. japonica (i.e., the mechanism by which tendrils of C. japonica avoid coiling around the stems of self plants) remains unclear.

Since self/non-self discrimination in C. japonica depends on the physiological connection [16], we predicted that electrical signals would enable self/non-self discrimination, just as in animals. To test this prediction, we measured the electrical potentials when tendrils contacted stems, which were connected via rhizomes (i.e., self plants). When the tendril did not coil around the stem, the electrical signal was generated from the stem to the tendril; when the tendril coiled around the stem, no electrical signal was generated. Based on this finding, we predicted that the electrical signals generated by the contact stimuli of the tendril inhibited the tendrils’ coiling. When the tendril contacted the stem and did not coil around the stem, it was removed from the stem and placed on a stick; the tendril coiled around the stick. Next, the part of the stem where the tendril did not coil was stimulated by contact. Then, the electrical signal was generated from the stem and transmitted to the tendril, causing the tendril to uncoil. The results revealed that the electrical signals generated by the contact stimuli inhibited the coiling of the tendril, and as a result, self/non-self discrimination was realised.

## 2. Results

We conducted touch experiments with transplanted C. japonica and detected electrical signals generated by tendrils’ contact (Fig. 1). Target plant tendrils were touched with stems of neighbouring plants. Neighbouring plants were connected via the rhizomes, and this type of connection is referred to as “RHIZOME” in ref. [16]. Electrical potentials were monitored with Ag/AgCl electrodes inserted into the following positions: stems with tendrils (the electrode is denoted by “Tendril”), stems of neighbouring plants (the electrode is denoted by “Stem”), and rhizomes (the electrode is denoted by “Rhizome”).

**Figure 1.**
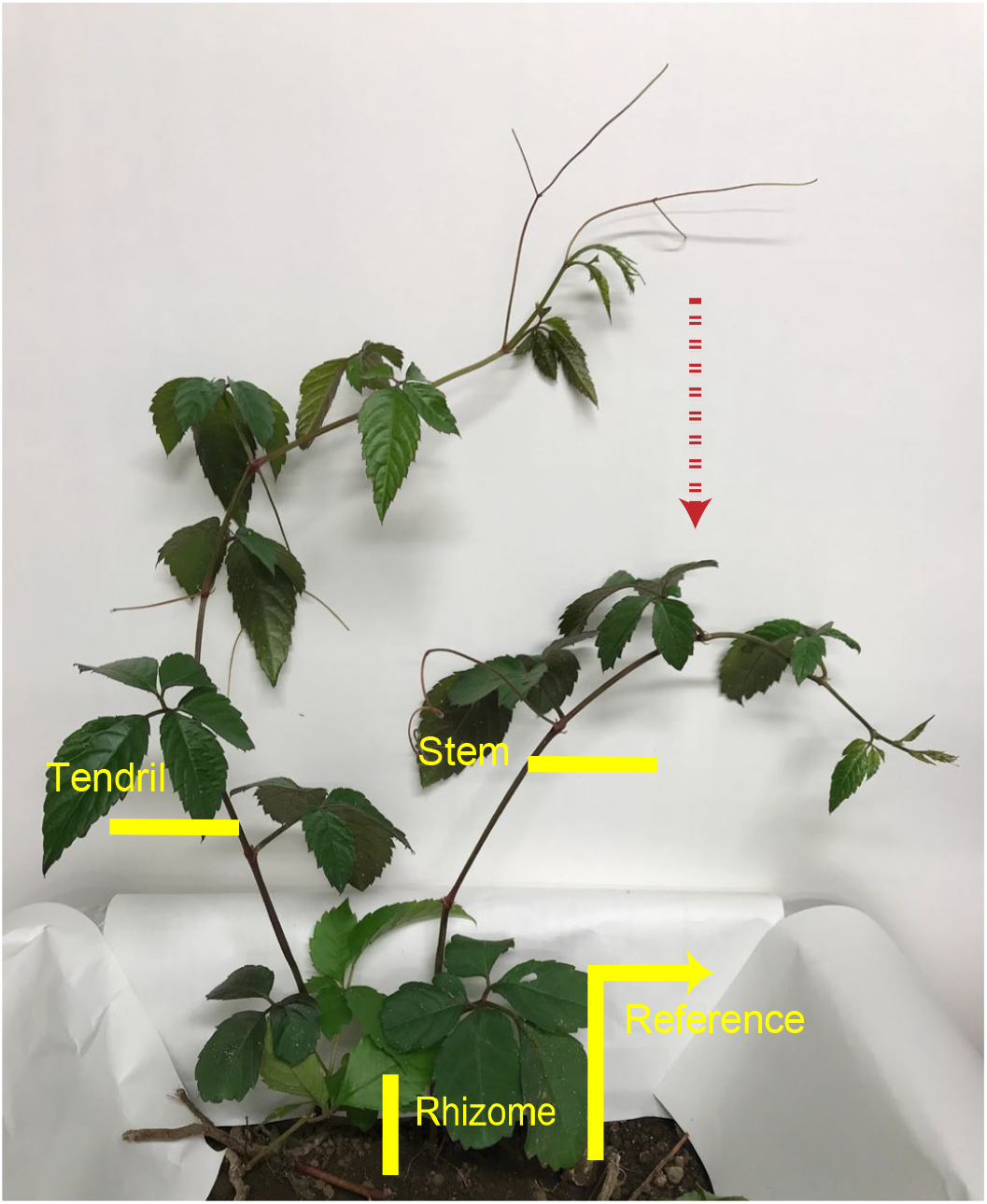
Electrode placement. Tendrils were brought into contact with stems of neighbouring plants. Electrical potentials were measured at electrodes on stems with tendrils (denoted by “electrical potential at the Tendril”), on stems where tendrils came into contact (denoted by “electrical potential at the Stem”), and on rhizomes (denoted by “electrical potential at the Rhizome”). The reference Ag/AgCl electrode was inserted into the soil.

### (a) Inhibition of tendril coiling with electrical signals

#### (I) Touch experiment involving the generation of electrical signals (Experiment 1)

The tendril was brought into contact with the stem (Fig. 2A). Fig. 3A shows the electrical potentials measured at each of the electrodes, Tendril, Stem, and Rhizome. When the tendril was in contact with the stem, the electrical potential at the Stem increased, followed by the electrical potential at the Rhizome, and then the electrical potential at the Tendril; that is, electrical signals were transmitted from the stem to the tendril via the rhizome. The electrical potential at the Tendril continued to increase, and the tendril did not coil around the stem. Next, the tendril was separated from the stem and brought into contact with the stick (Fig. 2B); then, the electrical potential at the Stem decreased. Similarly, the electrical potentials at the Rhizome and the Tendril also decreased. As a result, the tendril began to coil around the stick (Movie S1). This means that although the tendril could coil around the stick, it did not coil around the stem.

**Figure 2.**
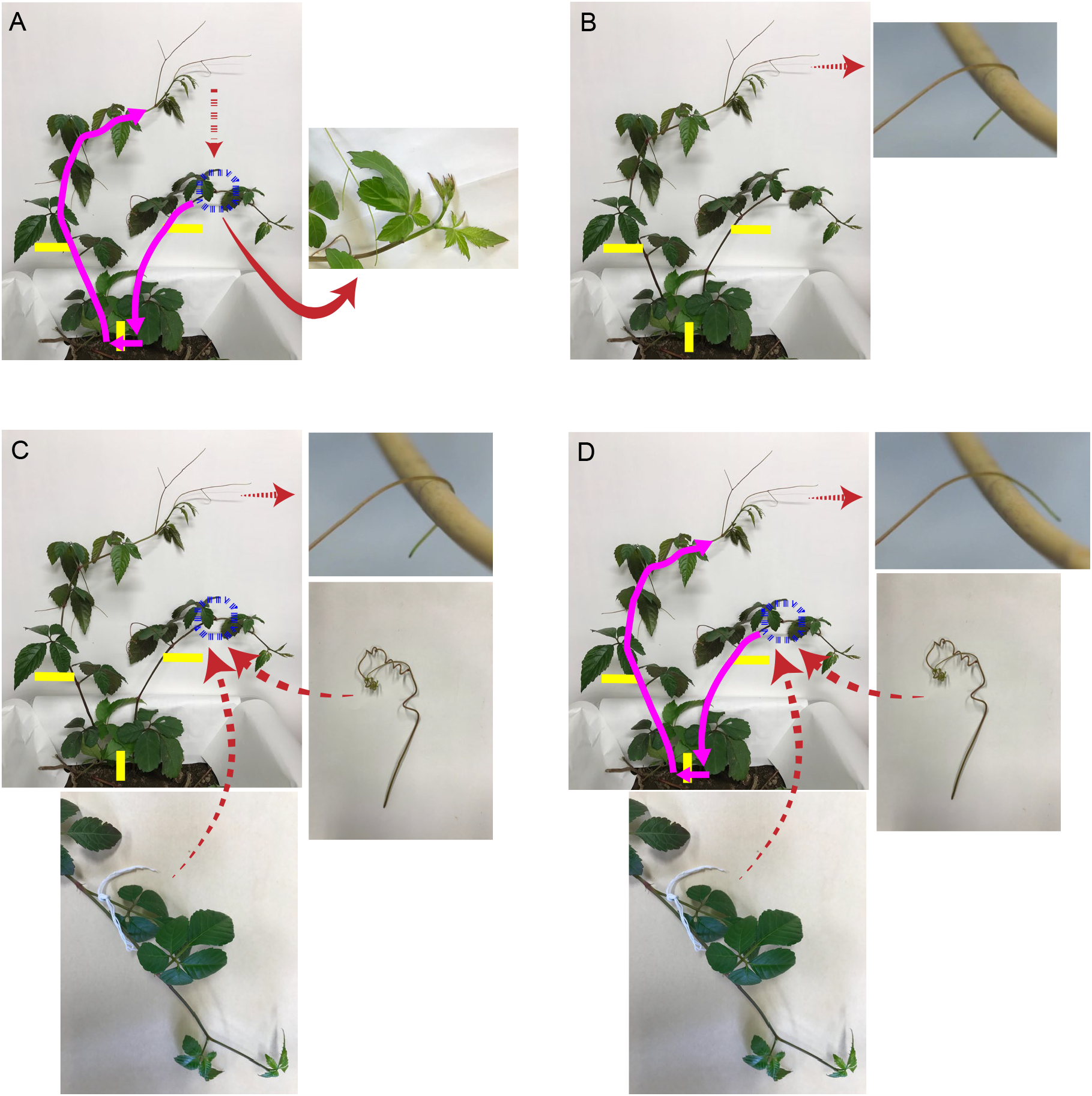
Illustration of experimental steps. (A) The tendril was touched with the stem of the neighbouring plant connected via the rhizomes. If the tendril began to coil around the stem, the tendril was brought into contact with other stems. When the tendril did not coil around, the electrical signal was generated from the position on the stem where the tendril made contact. (B) The tendril in contact with the stem was brought into contact with a stick; then the tendril coiled around the stick. (C) Two types of contact stimuli were applied to the position of the stem where the tendril had not coiled around in step A: (1) A tendril taken from another C. japonica was placed at that position. (2) A thread was tied to that position of the stem. (D) The contact stimuli in step C evoked the electrical signals, and the tendril uncoiled. The experiment was conducted with the C. japonica lying on its side so that the tendrils could easily contact the stems.

**Figure 3.**
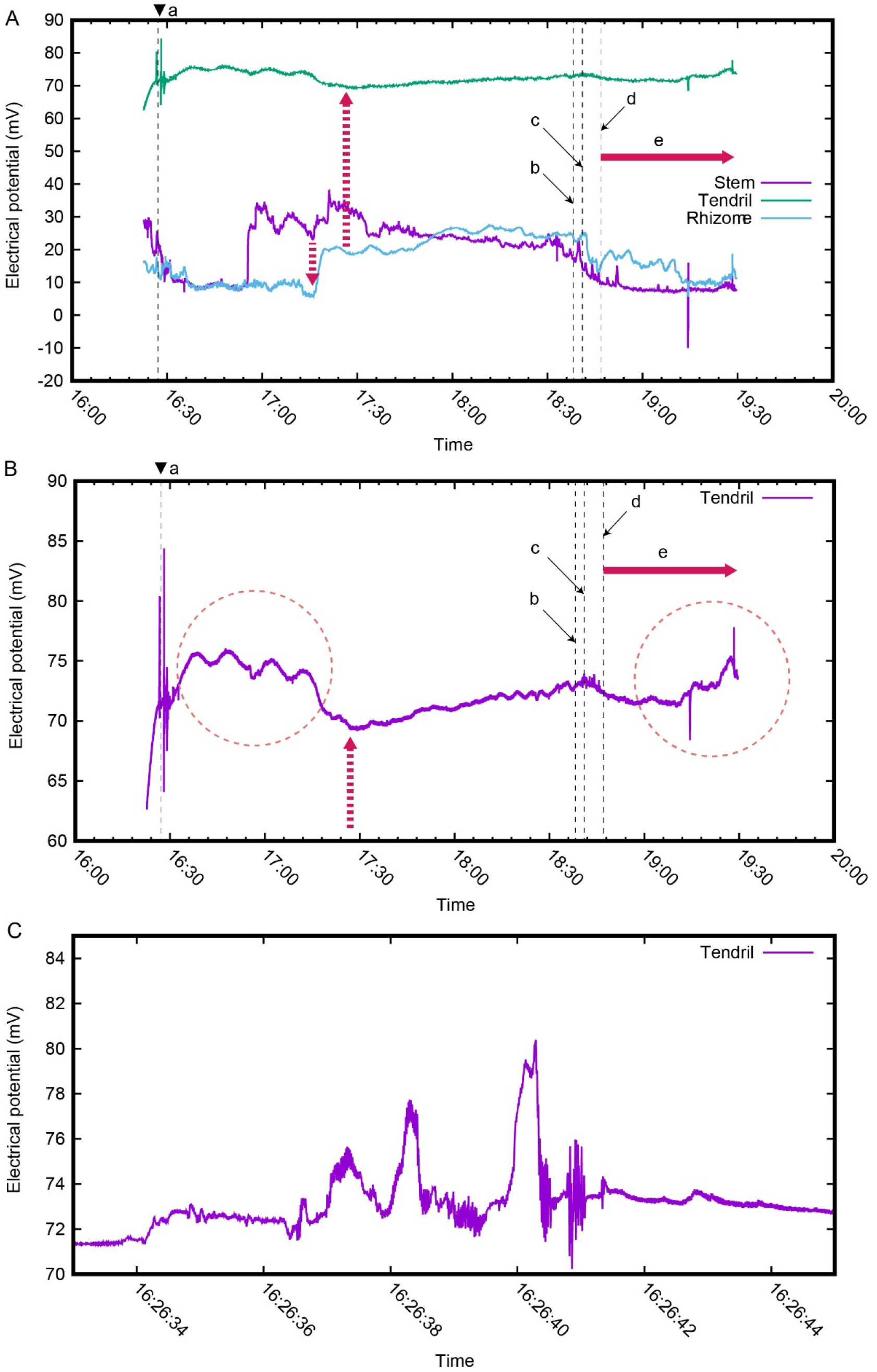
The electrical potential when the tendril did not coil around the stem. (A) The tendril was brought into contact with the stem (a). The tendril was removed from the stem because it did not coil around the stem (b). The tendril was brought into contact with the stick (c). The tendril began to coil around the stick (d). The tendril continued to coil around the stick (e). (B) The electrical potential at the Tendril. (C) The electrical potential at the Tendril was displayed in the range of 70 mV to 85 mV between 16:26:33 and 16:26:45.

In Figure 3B, the range of electrical potential was restricted to 60 mV to 90 mV, and only the electrical potential at the Tendril is shown. To bring the tendril into contact with the stem, the stem with the tendril was grasped by hand. During this period, there were contacts between the hand and the stem and between the tendril and the stem. These contacts generated the spike-like electrical potential (Fig. 3C). After the spike-like electrical potential generation, the electrical potential at the Tendril increased before the electrical signal arrived from the stem (the red circle on the left side of Fig. 3B). Similarly, the electrical potential at the Tendril increased when the tendril began to coil around the stick (the red circle on the right side of Fig. 3B).

#### (II) Touch experiment that did not involve the generation of electrical signals (Experiment 2)

After completing Experiment 1, the electrode was left pricked. The next day, the tendril was again brought into contact with the position on the stem that had been contacted the previous day. Then the tendril coiled around the stem. Fig. 4A shows the electrical potentials measured at each electrode. There was no increase in the electrical potential at the Stem with which the tendril was in contact, and there was no transmission of electrical signals from the stem to the tendril via the rhizome. As in the previous day’s experiment, when the tendril was brought into contact with the stem, the spike-like electrical potential was generated, and the electrical potential also increased (the red circle on the left side of Fig. 4B). The electrical potential then decreased, and the tendril began to coil around the stem. When the tendril perfectly coiled around the stem, the electrical potential increased again (the red circle on the right side of Fig. 4B) (Movie S2).

**Figure 4.**
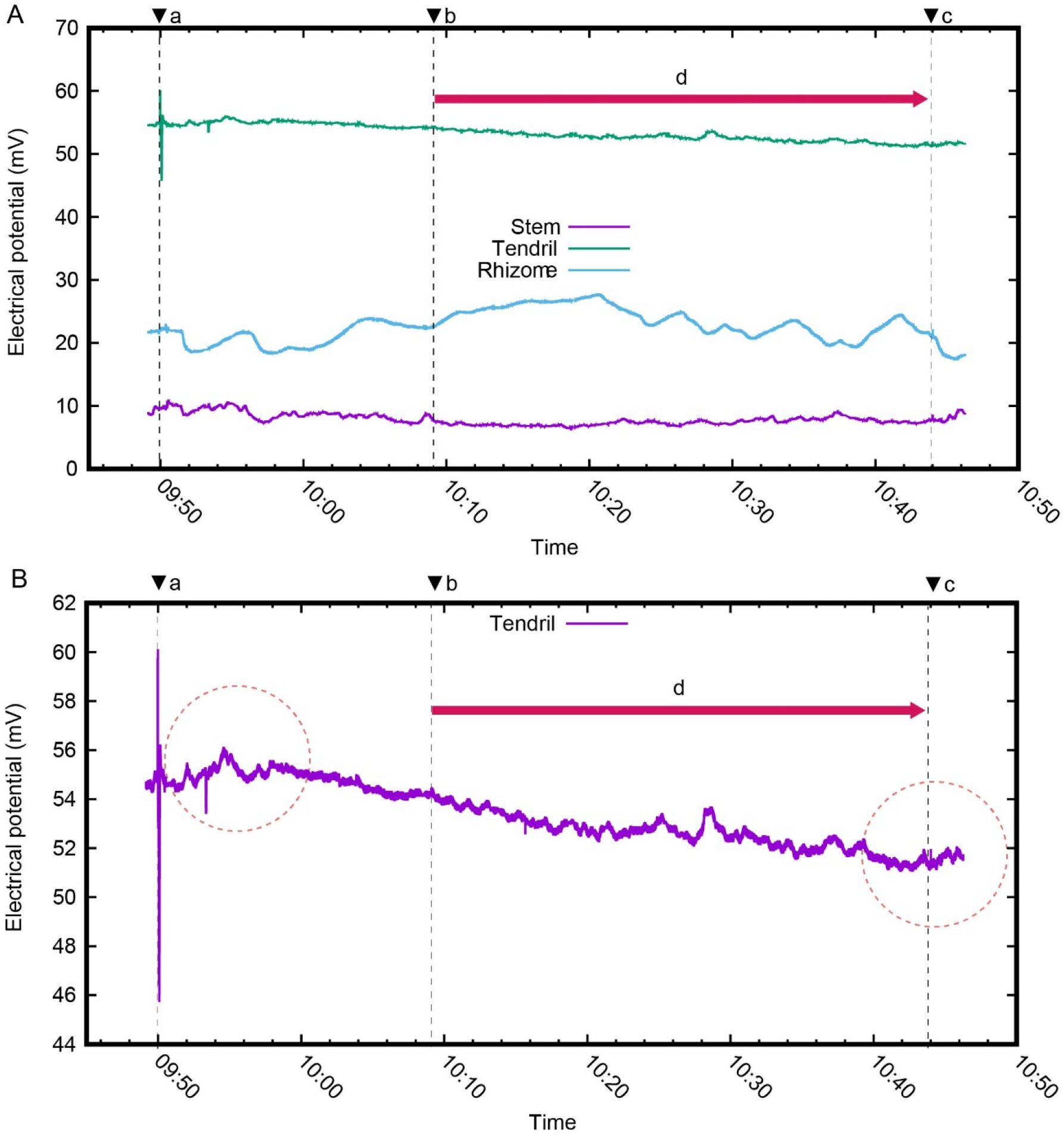
The electrical potential when the tendril coiled around the stem. (A) The tendril was brought into contact with the stem (a). The tendril began to coil around the stick (b). The tendril coiled around the stem perfectly (c). The tendril continued to coil around the stick (d). (B) The electrical potential at the Tendril was displayed in the range of 44 mV to 62 mV.

### (b) Inhibition of tendril coiling with electrical signals evoked by contact stimuli

Comparison of the results of Experiments 1 and 2 suggested that the electrical signal inhibited the tendril’s coiling. To confirm the inhibition by electrical signals, we artificially applied contact stimuli to the stem, generated electrical signals, and confirmed whether inhibition of the tendril’s coiling occurred. The experimental procedure was as follows:

1. The tendril was brought into contact with the stem (Fig. 2A).
2. The tendril was separated from the stem after making sure that the tendril did not coil around the stem.
3. The tendril was brought into contact with the stick (Fig. 2B).
4. The tendril began to coil around the stick.
5. Two types of contact stimuli were applied to the position of the stem where the tendril had not coiled around (Fig. 2C).
6. 6. The electrical signal evoked by the contact stimuli uncoiled the tendril. The experimental results when two types of contact stimuli were applied in step 5 above are shown below.

#### (I) Experiment stimulated by the tendril taken from another C. japonica (Experiment 3)

The electrical potentials when the stem was in contact with the tendril taken from another C. japonica are shown in Fig. 5A. When the tendril was brought into contact with the stem, the electrical potentials at the Stem, Rhizome, and Tendril all increased, and the tendril did not coil around the stem. However, because of the intermittent increase, the transmission of the electrical signal from the stem to the tendril was unclear.

**Figure 5.**
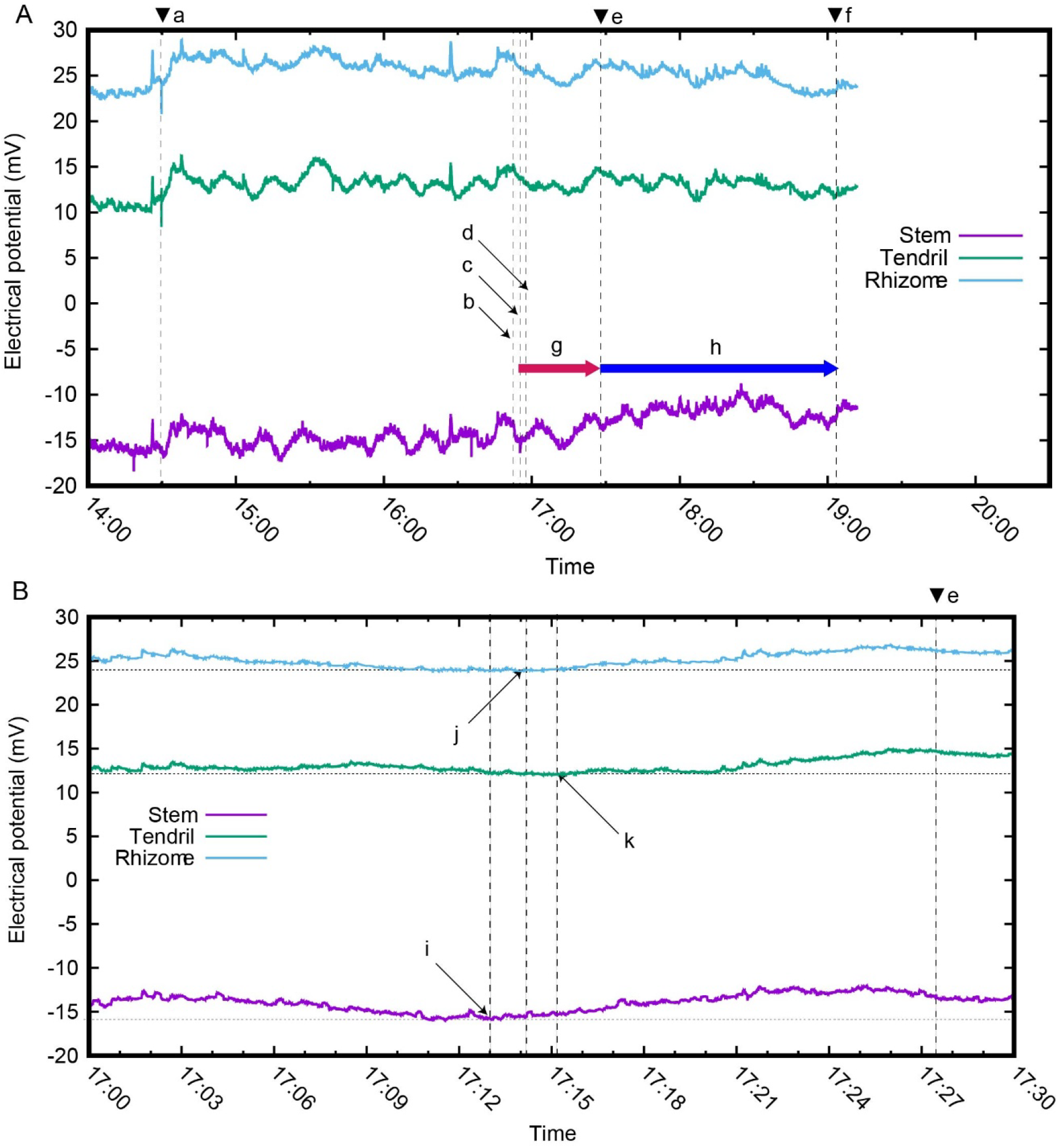
The electrical potentials evoked by the contact stimulus by the tendril taken from another C. japonica. (A) The tendril was brought into contact with the stem (a). The tendril was brought into contact with the stick (b). The tendril began to coil around the stick (c). The tendril taken from another C. japonica was placed at the position of the stem where the tendril had not coiled around (d). The tendril began to uncoil (e). The tendril uncoiled perfectly (f). The tendril continued to coil around the stick (g). The tendril continued to uncoil (h). (B) The electrical potentials between 17:00 and 17:30 were displayed. The electrical potential at the Stem increased (i). The electrical potential at the Rhizome increased (j). The electrical potential at the Tendril increased (k).

Although the signal transmission was unclear, the position of the stem where the tendrils did not coil around was identified. The tendril was removed from the stem and brought into contact with the stick. The electrical potential at the Tendril decreased, and the tendril began to coil around the stick. Next, when the tendril taken from another C. japonica was then placed on the stem where the tendrils did not coil around, the electrical potential at the Stem increased first, followed by the Rhizome and the Tendril (Fig. 5B), indicating that the electrical signal was transmitted from the stem to the tendril. As the electrical potential at the Tendril increased, the tendril began to uncoil (Fig. 5A, e) (Movie S3). The increase in electrical potential at the Stem when the tendril taken from another C. japonica was placed on the stem was greater than that increase when the tendril was in contact with the stem (Fig. 5A).

#### (II) Experiment stimulated by the thread (Experiment 4)

The electrical potentials are shown in Fig. 6 when the contact stimulus was applied by tying the thread to the stem instead of the tendril taken from another C. japonica. When the tendril was brought into contact with the stem, the electrical signal was transmitted from the stem to the tendril via the rhizome (Fig. 6B), and the tendril did not coil around the stem. When the tendril was brought into contact with the stick, the tendril coiled around the stick; when the position of the stem, where the tendril did not coil around, was tied with a thread tightly, the electrical potential at the Tendril increased, and the tendril uncoiled, as in Experiment 3. Since, after the tendril came into contact with the stick, the electrical potentials at the Stem, the Rhizome, and the Tendril remained nearly constant, the transmission of the electrical signal was unclear (see Discussion). After the tendril left the stick, the electrical potentials of these three electrodes increased rapidly (Fig. 6A, 6C). Comparing the times at which the electrical potentials increased shows that the electrical signal was transmitted from the stem to the tendril (Fig. 6D).

**Figure 6.**
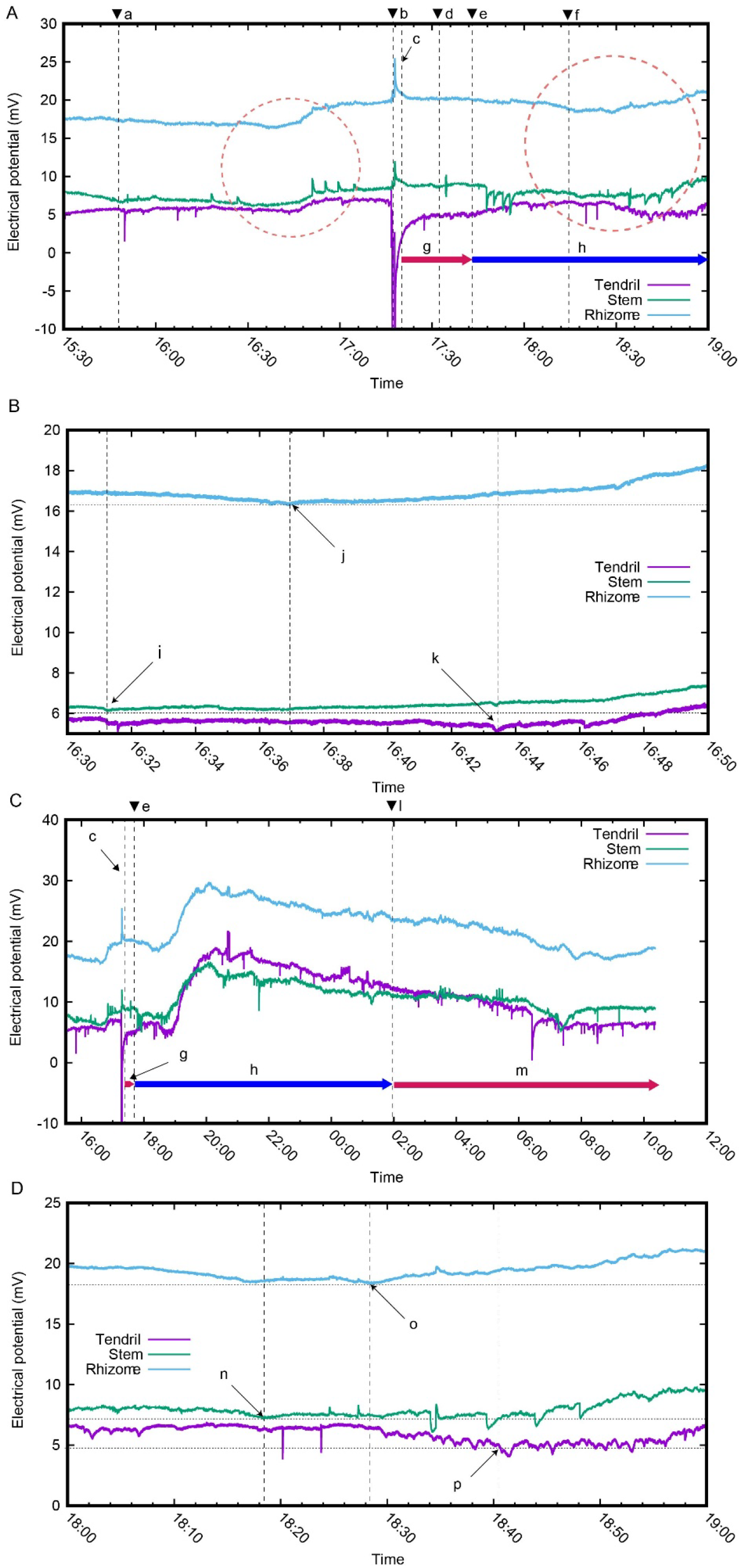
The electrical potential when the contact stimulus was applied by tying the thread to the stem. (A) The tendril was brought into contact with the stem (a). The tendril was removed from the stem and brought into contact with the stick. (b) The tendril began to coil around the stick (c). The stem was tied tightly with the thread (d). The tendril began to uncoil (e). The tendril uncoiled perfectly (f). The tendril continued to coil around the stick (g). The tendril continued to uncoil (h). A spike-like electrical potential at the Tendril of about -70 mV was generated when the tendril was in contact with the stick. To enhance the visibility of the electrical potential change, the electrical potentials were displayed in the range of -10 mV or higher. (B) The electrical potentials were displayed in the range of 5 mV to 20 mV between 16:30 and 16:50 (the red circle area on the left side of Fig. 6A). The electrical potential at the Stem increased (i). The electrical potential at the Rhizome increased (j). The electrical potential at the Tendril increased (k). (C) The tendril began to coil around the stick again (l). The tendril continued to coil around the stick (m). (D) The electrical potentials were displayed in the range of 0 mV to 25 mV between 18:00 and 19:00 (the red circle area on the right side of Fig. 6A). Spike-like electrical potentials were generated at the Stem and the Tendril. By ignoring these spike-like electrical potentials, the time of the electrical potential increase was determined. The electrical potential at the Stem increased (n). The electrical potential at the Rhizome increased (o). The electrical potential at the Tendril increased (p).

The tendril movement was observed overnight after the tendril had separated from the stick. The changes in the electrical potentials until the following day are shown in Fig. 6C. The electrical potentials of the three electrodes increased rapidly, then decreased gradually; the tendril again coiled around the stick (Fig. 6C, Movie S4).

### (c) Velocities of electrical signals

Five contact stimuli were applied to the stems of C. japonica plants, and velocities of electrical signals were measured. The five measurements of the velocity of the electrical signal were 0.11, 0.11, 0.11, 0.5, and 0.78 cm/s.

## 3. Discussion

We investigated the electrical signals generated when the tendrils came into contact with the stems. By comparing the results from experiments 1 and 2, we predicted that the electrical signals generated by the contact stimuli of the tendril inhibited the tendrils’ coiling. Thus, two types of objects (the tendril taken from another C. japonica and the thread) were brought into contact with the stem to test (1) whether electrical signals could only be generated by contact stimuli by the tendrils (or whether electrical signals were also generated by contact stimulus by the thread), and (2) whether these generated electrical signals inhibited the tendrils’ coiling. The results of Experiments 3 and 4 showed that not only physiologically connected tendrils but also contact stimuli by other substances generated electrical signals, and that these electrical signals inhibited the coiling of tendrils. Furthermore, the results of Experiment 2 indicated that electrical signals were not always generated by contact stimuli; when they were not generated, the tendrils coiled around the stems of self plants (see legend of Movie S4).

The increase in electrical potential at the Stem (approximately 2 mV) that occurred when the tendril was brought into contact with the stem was much smaller than the increase in electrical potential at the Stem (approximately 8 mV) when the same position on the stem was tied tightly with the thread (Fig. 6C). This means that the electrical potential varies with the intensity of the contact stimuli, which is consistent with the characteristics of VPs [13]. The velocity of the electrical signal was in the range 0.11-0.78 cm/s, which is also consistent with the range of VPs (0.1–1.0 cm/s) [12]. These results identify the electrical signals generated by the contact stimuli as VPs. One of the roles of VPs in plants has been found to activate defences in tissues distal to wounds [23]. Conversely, APs regulate rapid leaf movements in touch-sensitive plants such as Mimosa and Venus flytrap, mediating the bending of sundew leaf tentacles [12]. Previous studies have reported that contact stimuli generate APs in tendrils [7, 8]. In the present study, contact stimuli generated the VP with AP-like spikes [24] and subsequently the VP (Fig. 3). The stimuli that evoke APs and VPs are unclear [25]. We found a new role for VPs in plants: VPs inhibited the coiling of the tendrils.

In Experiment 4, after the tendril was separated from the stem and brought into contact with the stick, the contact stimulus to the stem was no longer present; therefore, the electrical potential at the Stem was predicted to decrease (see Fig. 6A). However, the electrical potential at the Stem remained constant because the threads were tied tightly (Fig.6A, d). This applied a strong contact stimulus, causing an increase in electrical potential sufficient to compensate for the decrease in electrical potential. The increase in electrical potential was transmitted to the rhizome and the tendril, and the electrical potential at the Tendril increased, which seems to have caused the uncoiling of the tendril. The transmission of electrical signals could not be confirmed because no increase or decrease in electrical potential occurred in the stem, the rhizome, and the tendril. On the other hand, Fig. 6D shows that the subsequent electrical signal (a significant increase in electrical potential after uncoiling) was transmitted from the stem to the tendril.

Although tendrils are more likely to coil around neighbours connected via rhizomes than neighbours connected via stems [16], we selected plants connected via the rhizomes because the presence of rhizomes made it easier to observe the transmission due to the time difference in the arrival of electrical signals at the three electrodes: Stem, Rhizome, and Tendril. In Experiment 1, after the tendril came into contact with the stem, the electrical potential at the Tendril increased before the electrical signal arrived from the stem (indicated by the red circle on the left side of Fig. 3B). Because the rhizomes of this experiment were long, it took time for the electrical signal to reach the Tendril from the Stem. The fact that the electrical potential at the Tendril increased before the arrival of this electrical signal shows that the tendril also generated the electrical signal when the tendril came into contact with the stem. Similarly, the tendril generated the electrical signal when it began to coil around the stick (indicated by the red circle on the right side of Fig. 3B).

A difference in the electrical potential at the Tendril was observed when the tendril first coiled around the stick (about 5 mV) and when it coiled around the stick a second time (about 10 mV) (Fig. 6C). Comparing the relationship between electrical potentials and coilings of the tendrils in Figures 3 through 6 implies that the tendrils begin to coil when the electrical potential at the Tendril decreases or when the electrical potential reaches a specific range, but further investigation is needed.

At present, the stimulus transduction from contact to coiling remains unclear. Jaffe revealed that both ventrally stimulated coiling and dorsally stimulated inhibition of coiling can be temporarily stopped by a 9-minute cold break at 10 °C, given immediately after stimulation [26]. It is unclear whether the VPs, as well as the cold break, inhibit the same event in the process of stimulus transduction. Further study of VP inhibition of coiling may help to understand the mechanism of the tendril’s coiling.

## 4. Materials and Methods

### (a) Plant Material and Growth Conditions

All C. japonica rhizomes used in this study were collected from the University of Electro-Communications. The plant is widespread in Japan and is neither a protected nor an endangered species. We selected neighbouring plants connected via the rhizomes, with at least one tendril. Rhizomes were cut into approximately 30 cm lengths and transplanted into flowerpots filled with soil collected from the University of Electro-Communications. Plants were planted horizontally so that the tendrils could contact the stems easily in the experiments. The plants were watered sufficiently. The experiments were conducted indoors at a temperature of about 25 °C.

### (b) Ag/AgCl electrodes

The Ag/AgCl electrodes consisted of pieces of silver wires (99.99% purity; A-M Systems, Inc., cat no. 788000) with a diameter of 0.762 mm, which were chlorinated and coated with an AgCl film. The length of the silver wire was variable according to the thickness of the stem or rhizome where it was inserted. To measure stable electrical potentials, measurements were taken at least 3 hours after the electrodes were inserted [27, 28]. The vascular bundles are arranged in a ring because C. japonica is a dicotyledonous plant. It is difficult to know in advance where the electrical signals pass through the vascular bundles in a ring during the tendril’s contact. If too many electrodes were inserted, C. japonica wilted; therefore, two electrodes were inserted into the stems, the rhizomes, and the stems with tendrils at a distance and in the orthogonal direction. As a result, the amount of change in the electric potential at the Stem, the Rhizome, and the Tendril did not always coincide because the distances between each electrode and the vascular bundles to which the electrical signal was transmitted could not be made equal.

Since electrical signals as derived from a number of environmental stimuli can travel at high speed over long distances throughout the entire plant [12], the observed electrical signals are complex. Of the two electrical potentials measured at each site, we selected the electrical potential that fluctuated closer to the timing of the contact stimuli and the movements of the tendril.

### (c) Measurement system

Figure 7 shows the schematic diagram of the measurement system. Low-level voltages were measured using probes—silver wire needles—attached to C. japonica. These signals were transmitted to a data logger (Hioki MR8875 Memory Hicorder) via coaxial cables and recorded as digital data. To measure the spike-like electrical potential, data were captured at a 10 ms interval (10 ms recording interval). The recorded data was subsequently transferred to a personal computer (PC) for further processing.

**Figure 7.**
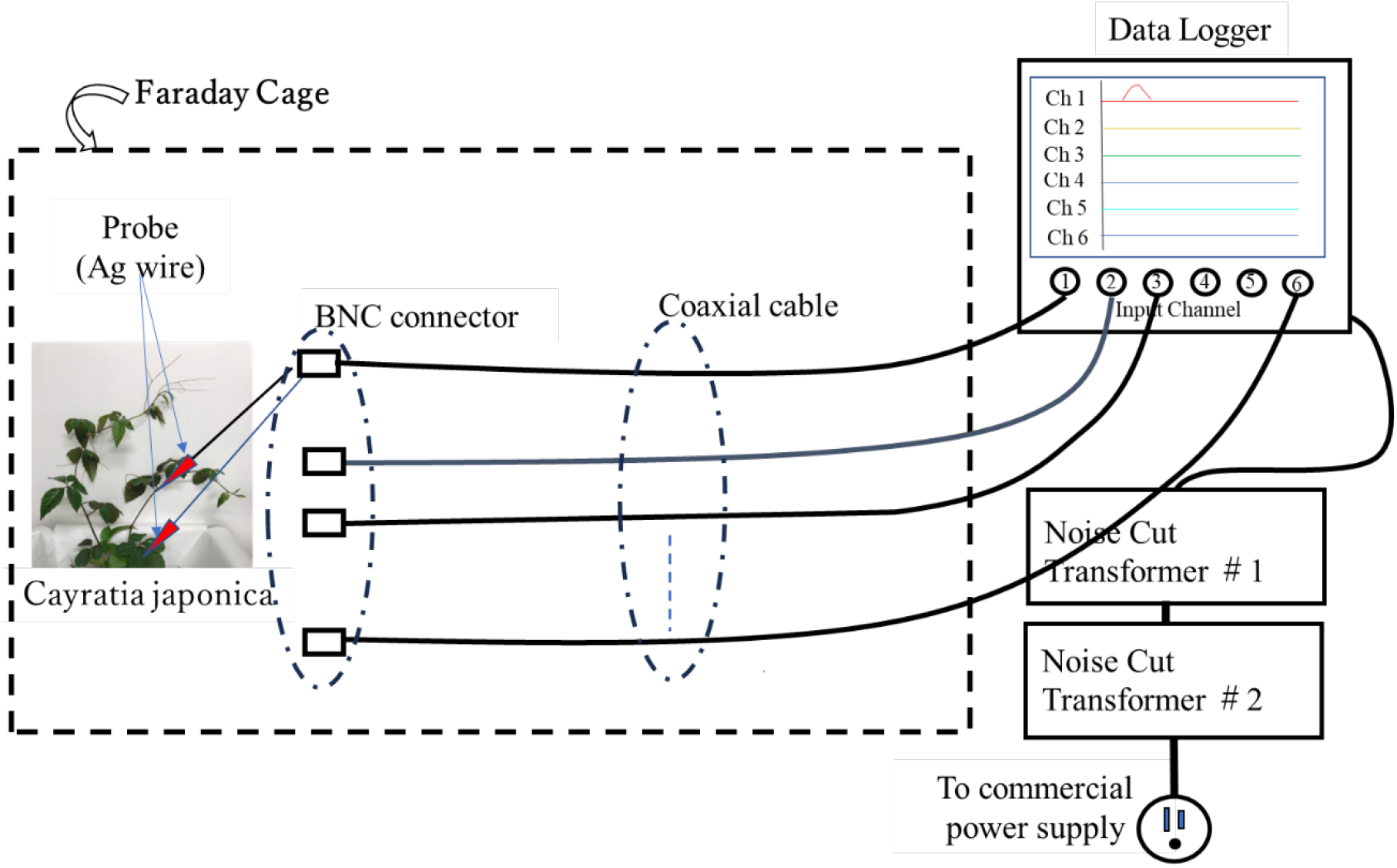
Schematic diagram of the measurement system: one set of probes is shown.

In general, electronic devices must neither emit nor be susceptible to electromagnetic (EM) noise. A device that satisfies both conditions is said to exhibit electromagnetic compatibility (EMC) [29]. The following section describes the measurement system incorporating EMC countermeasures tailored for low-level voltage measurements. Special attention is given to minimising interference from external EM noise, particularly in the measurement probes and circuits. Electromagnetic (EM) noise, a type of EM wave, comprises electric and magnetic fields. These fields can induce currents in sensitive circuits, significantly affecting low-level voltage measurements. Additionally, broadcast and communication signals may act as unwanted noise for unrelated devices. To minimise EM noise intrusion, the following countermeasures have been implemented in the measurement system.

### (d) Radiated Interference Countermeasures

External electromagnetic (EM) noise can induce unwanted currents in measurement probes and associated wiring, thereby distorting the measured voltage signals. In this system, the voltages obtained from the probes were transmitted to a data logger via coaxial cables with high shielding effectiveness. To minimise the influence of external EM fields on the wiring connected to the probes and the transmission lines, various shielding and layout strategies have been implemented. A Faraday cage was employed to reduce the impact of external EM fields, and all measurements were conducted within its enclosed space. The cage was constructed using a metal mesh. When the frequency of the EM noise to be suppressed is sufficiently low (i.e., long wavelengths), the openings in the mesh become small relative to the wavelength, enabling the cage to shield against radiated interference effectively. The Faraday cage used in this measurement system was a cuboid with a base measuring 1.2 m × 0.9 m and a height of 0.9 m. It was made of a honeycomb metal mesh with cell sizes of approximately 1 cm. The voltage waveforms obtained from C. japonica exhibited gradual rise times in the millisecond range in the time domain, indicating that their frequency components were very low and their wavelengths were extremely long. Consequently, the mesh openings were negligible compared to the wavelength of the signals of interest. Even radio signals at much higher frequencies, such as 540 kHz (wavelength ≈ 560 m), were difficult to receive inside the cage, confirming that the targeted EM waves were effectively shielded.

### (e) Conducted Interference Countermeasures

The data logger used in this system required a 50 Hz sine wave AC power supply, which was provided by a commercial power source. However, various devices connected to the power grid generate noise that propagates through the power lines. This noise, classified as conductive interference, must be effectively suppressed to ensure accurate measurement. To address this issue, a power filter was employed to eliminate noise entering through the commercial power supply. The filter consists of a high-performance noise-cut transformer designed for effective noise suppression. To further enhance its filtering capability, two such transformers were connected in cascade, reducing the noise voltage to approximately 1/10000 of its original value. As a result, impulse-like noise caused by the operation of nearby elevators and conductive noise from digital devices and personal computers were attenuated to negligible levels. By performing the countermeasures, this system enabled stable and reliable measurement of the low-level voltages generated by C. japonica.

## Acknowledgments

T.H. thanks Yuki Homma for editing movies.

## Supporting Information

### Movie S1 (separate file)

Video of the tendril’s contact with the stem and the stick. The tendril did not coil around the stem, but it coiled around the stick. The stick tilted slightly while the tendril was coiling around the stick due to an incomplete fixing. The experiment was stopped before the tendril completely coiled around the stick. Once the tendril completely coils around the stick, the experiment cannot be conducted again. By aborting the experiment, as a result, Experiment 2 could be conducted the next day.

### Movie S2 (separate file)

Video of the tendril’s contact with the stem. The tendril was contacted again at the position on the stem where it had been contacted the previous day.

### Movie S3 (separate file)

Video of contact stimulus by the tendril taken from another C. japonica.

### Movie S4 (separate file)

Video of contact stimulus by tying the thread to the stem. Since the tendril coiled around the first stem with which it was brought into contact, it was brought into contact with the neighboring stem again. The image of the tendril initially coiled around the stem is omitted. Since the tendril had coiled around the first stem, it was already bent when it came into contact with the next stem. Two tendrils were growing from a single stem. After the tendril coiled around the stick again, the other tendril also coiled around the stick.

